# Point Process Temporal Structure Characterizes Electrodermal Activity

**DOI:** 10.1101/2020.03.11.982843

**Authors:** Sandya Subramanian, Riccardo Barbieri, Emery N. Brown

## Abstract

Electrodermal activity (EDA) is a read-out of the body’s sympathetic nervous system measured as sweat-induced changes in the electrical conductance properties of the skin. There is growing interest in using EDA to track physiological conditions such as stress levels, sleep quality and emotional states. Standardized EDA data analysis methods are readily available. However, none considers two established physiological features of EDA: 1) sympathetically mediated pulsatile changes in skin sweat measured as EDA resemble an integrate-and-fire process; 2) inter-pulse interval times vary depending upon the local physiological state of the skin. Based on the anatomy and physiology that underlie feature 1, we postulate that inverse Gaussian probability models would accurately describe EDA inter-pulse intervals. Given feature 2, we postulate that under fluctuating local physiological states, the inter-pulse intervals would follow mixtures of inverse Gaussian models, that can be represented as lognormal models if the conditions favor longer intervals (heavy tails) or by gamma models if the conditions favor shorter intervals (light tails). To assess the validity of these probability models we recorded and analyzed EDA measurements in 11 healthy volunteers during 1 to 2 hours of quiet wakefulness. We assess the tail behavior of the probability models by computing their settling rates. All data series were accurately described by one or more of the models: two by inverse Gaussian models; five by lognormal models and three by gamma models. These probability models suggest a highly succinct point process framework for real-time tracking of sympathetically-mediated changes in physiological state.

## Introduction

Electrodermal activity (EDA) is the most common definition used to describe changes in the electrical properties of the skin in response to a person’s physiological or emotional state as captured by the sympathetic nervous system (1). At all times, there is a background state of EDA due to some filling of sweat glands. Changes in this level of filling occur in response to internal stimuli (physiological and emotional) and external stimuli such as threats. EDA measures the activity of eccrine sweat glands in the dermis and epidermis in response to changes in sympathetic activity. This is one of the most primal components of the flight-or-fight response. Therefore, EDA serves as a measure of the sympathetic nervous system activity in clinical studies such as lie detector tests and assessment of stress (1). It is also being developed as a neuromarketing tool to evaluate consumer responses to different products or promotions. For this reason, there is growing interest in the development of real-time algorithms to accurately characterize EDA.

EDA shows a characteristic pattern. First, two distinct levels of activity occur simultaneously. There is a baseline or tonic component that drifts gradually across time. Second, on top of this component is the phasic component which consist of pulse events that vary dramatically in amplitude, shape, width, and spacing. The phasic component is believed to respond more quickly to fast-changing sympathetic nervous system activity. Third, there is a relationship between the EDA baseline and the inter-pulse intervals as well as between the EDA baseline and the pulse amplitudes. Fourth, there is a large variation in both baseline activity and pulse activity within subjects and between subjects.

Most current EDA analysis methods focus mainly on the phasic component to characterize sympathetic activity. These methods fall primarily in two categories: rate-based methods and deconvolution methods. The rate-based methods specify a particular time window and provide as output the number of pulse events per time window (2–4). For the deconvolution methods, a single pulse shape is assumed for each subject, and the EDA signal is represented as the convolution of this pulse shape with neural inputs. Therefore, the neural input is deconvolved from the EDA while also simultaneously fitting the optimal pulse shape parameters (5–9). Hence, the deconvolution methods report the occurrence times and amplitudes of the pulse events.

While the rate-based and deconvolution methods are widely used, they have important shortcomings. Most importantly, they do not use the well-established physiology of EDA in their construction. Also, formal statistical modeling is not used to characterize the inter-pulse interval dynamics (pulse rate and pulse times) or the pulse amplitudes.

The important advances that we report are represented by point process models derived from EDA physiology to characterize inter-pulse interval dynamics. These mathematical constructs provide a novel, physiologically-based frame work for EDA analysis. The balance of this paper is organized as follows. In Physiology and Theory, we use the well-known features of EDA physiology to formulate statistical models of EDA. Then in Application and Results, we illustrate use of these models in the analysis of EDA recordings from eleven healthy subjects during the awake-state and during rest. Finally, in the Discussion, we summarize the models and the implications.

## Physiology and Theory

### The Anatomy and Physiology of Electrodermal Activity

To develop our statistical models of EDA activity we review the anatomy and physiology of sweat production in the skin. Each eccrine sweat gland consists of three parts, the dermal gland, the duct that connects the gland to the skin surface, and the pore where the duct opens to the skin. The dermal portion of the gland is innervated by sudomotor nerves, a part of the peripheral nervous system that is predominantly under sympathetic control. Sweat is produced in the gland by sympathetically-induced abrupt increases in spiking activity in the sudomotor nerves. These pulsatile events are called sudomotor bursts. Sweat produced in response to these bursts accumulates in the duct where it reaches the skin surface by pushing open the pore. At the same time, sweat dissipates by constant reabsorption through the walls of the duct and by evaporation from the skin surface (9).

Sweat on the skin increases the skin’s electrical conductance (inverse of resistance) because the salt containing sweat “completes the circuit”. The electrical conductance across the skin can be measured in a standard fashion by placing two electrodes on either the palm or fingers, applying a constant voltage, and measuring the current. The pulsatile effects of the sudomotor bursts measured at the skin are termed galvanic skin responses (GSRs). The second-to-second changes in skin sweating measured as second-to-second changes in skin conductance is termed EDA.

EDA measurements have two distinct components. First, there is background or tonic activity in both sudomotor nerve bursts and in EDA in the absence of specific sympathetic stimuli (10–15), but bursts of sudomotor nerve activity do not always translate one-to-one into GSR events. Background GSR events are typically small in amplitude and have large inter-event gaps. Studies of sudomotor nerve activity have established that background EDA is characterized by this lack of a one-to-one relationship. This relationship is highly dependent upon ambient conditions and determines the baseline filling state of the sweat gland. For example, tonic activity and gland filling are higher when the skin temperature is warm and lower when the skin temperature is cold.

The second component of EDA measurements are the frequency and intensity of sympathetic-induced stimulation. The baseline level of sweat gland filling modulates the ease with which any sudomotor nerve activity (spontaneous or stimulated) induces a conductance change, and the size of that change (Figure S1). The greater the proportion of sweat production required to fill the gland initially, the smaller the resulting pulse amplitude. Most importantly, the studies show that that the GSR amplitude correlates more strongly with the background physiologic state than with the magnitude of the sympathetic stimulus. Hence, despite maintaining constant stimulus intensity, GSR amplitudes can increase purely based on increases in the tonic level. In fact, after a certain point, continued increases in the background state actually decrease GSR amplitudes even though stimulus intensity is held constant (14–19). Therefore, relying on GSR amplitudes as measures of stimulus intensity, especially in the presence of a dynamic background, can be misleading (10–27).

### Statistical Model of Galvanic Skin Responses

We will investigate four classes of statistical models to describe the time series of GSR events recorded during EDA measurements: inverse Gaussian, exponential, gamma and lognormal. This choice of models follows directly from the physiology. We first assume a stable background level of sympathetic activity leading to a baseline level of sweat in the duct. Assume that the production of sweat in the gland and its release onto the skin in response to sympathetic stimulation, obeys an integrate-and-fire model defined by a Gaussian random walk with drift commonly used in computational neuroscience. The Gaussian random walk with drift describes the time course of sweat accumulation in the duct in response to sympathetic stimulation. More stimulation leads to greater sweat accumulation depicted by the random walk integrating or moving in a positive direction. Sufficient sweat accumulation in the duct pushes open the pore producing an increase in skin conductance measured as a discrete GSR event (1). This corresponds to the random walk crossing a threshold and the neuron firing. The neuron resets and process can start again. Similarly, the duct empties through evaporation and reabsorption and the accumulation process can begin again with the next round of sympathetic stimulation. It is well known that the distribution of times between threshold crossing events for the Gaussian random walk with drift process is the inverse Gaussian probability model (28,29). This model has been successfully used to describe a range of point process phenomena, including neural spiking activity, heartbeats, geyser eruptions and earthquake-after shock occurrences (30).

We postulate that under stable, or approximately stable, background conditions the inverse Gaussian model will provide an accurate description of the GSR inter-event times as the integrate and fire model is plausible description of skin sweat production. We postulate that under variable background conditions, there may be a mixture of inverse Gaussian models. This could produce either longer inter-event intervals or shorter inter-event intervals relative to a single inverse Gaussian model. The former case might occur with cold skin and hence, a low level of background activity. The latter case may occur with warm skin and a higher level of background activity. The low background activity would result in heavier tails of the distribution whereas the higher background activity would result in lighter tails of the probability distributions. To model heavy tail distributions, we consider the lognormal and the gamma distributions; whereas to model the light tail distributions, we consider the gamma and the exponential distributions. Although the exponential distribution is a special case of the gamma, it represents the null model of a Poisson process underlying GSR event production. The gamma is the most flexible of the distributions and can have either a heavier or lighter tail than the inverse Gaussian depending on the values of the parameters.

We quantify the heaviness of the distribution tails by evaluating the asymptotic settling rate which is defined as

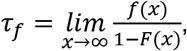

where *f*(*x*) is the inter-event probability model and *F*(*x*) is the corresponding cumulative distribution function.

Based on this definition, a distribution is classified as light, medium, or heavy tailed if the settling rate is infinity, positive but finite, or zero respectively. In other words, the slower the tail of a distribution “settles” or the lower the settling rate, the heavier the tail. Based on tail behavior alone, these four distributions can be divided into three medium-tailed (exponential, gamma, and inverse Gaussian) and one heavy-tailed distribution (lognormal). The three medium tailed distributions are distinguished based on their parameter values, which determine the respective settling rates (31,32).

## Application

### Experimental Data

With the approval of the Massachusetts Institute of Technology (MIT) Institutional Review Board, we collected EDA data from 12 healthy volunteers between the ages of 22 and 34 while awake and at rest. Electrodes were connected to the second most distal phalange of the second and fourth digits of each subject’s non-dominant hand. Approximately one hour of EDA data was collected at 256 Hz. Subjects were seated upright and instructed to remain awake. They were allowed to read, meditate, or watch something on a laptop or tablet, but not to write with the instrumented hand. One subject's data were not included in the analysis because they informed us after the data collection that they occasionally experience a Raynaud's type phenomenon. This would affect the quality of their EDA data. Data from the remaining 11 subjects were analyzed using Matlab 2017a. All data and code are available on the Neuroscience Statistics Research Laboratory website (www.neurostat.mit.edu).

### Data Preprocessing and Galvanic Skin Response Event Selection

Preprocessing consisted of two major steps, 1) detecting and removing artifacts and 2) isolating the phasic component. Artifact detection was done based on the derivative of the time series, since large rapid changes are physiologically impossible for skin conductance. Artifact removal was done in two parts, first correcting for artifact-related large magnitude changes in the remainder of the signal, and then interpolating the few seconds around the artifact itself. Then a low-pass FIR filter was used to estimate and remove the slow-moving tonic component of the signal, isolating the phasic component. The data preprocessing is described in further detail in Subramanian et al. (33).

Based on physiology, knowing the absolute pulse amplitude alone is insufficient to extract pulses reliably. Therefore, we computed locally-adjusted amplitudes for all detected peaks using the Matlab function *findpeaks*. The *findpeaks* algorithm computes a 'prominence' or relative amplitude for each peak, which adjusts the amplitude of each peak as the height above the highest of neighboring “valleys” on either side. The valleys are chosen based on the lowest point in the signal between the peak and the next intersection with the signal of equal height on either side. With this method, a peak with small absolute amplitude can be 'rewarded' in its prominence value if it is in a region of data with very low activity. We then used a threshold on this prominence value instead of the absolute amplitude, which automatically accounted for the dependence of pulse amplitude on background state: in low background states, pulses are smaller. Instead of dynamically changing thresholds, the algorithm adjusted relative amplitudes.

We used a prominence threshold of 0.005 to extract peaks across all subjects, unless this resulted in too few or too many pulses for each hour-long recording. This was primarily verified by visual inspection of extracted pulses, as well as rough estimates of 60 and 360 pulses as generous bounds for what should be expected of one hour of data at rest with little external stimulation. This corresponds to one pulse every 10-60 seconds on average (3,10–15). In the case too few pulses were extracted, we gradually reduced the prominence threshold by 0.001 until the number of pulses exceeded 100 and no obvious pulses were missed by visual inspection (verified by three different viewers independently). In the case too many pulses were extracted, we gradually increased the prominence threshold by 0.001 until the number of pulses was less than 360 and the extracted pulses did not look like they were picking up sensor noise by visual inspection (verified by three different viewers independently). If changing the threshold in increments of 0.001 resulted in drastically changed the number of pulses each time, we reduced the increment to 0.0005. Though not fully automated at this stage, the design of this method to extract pulses attempted to take into account the wide variation in baseline levels of EDA activity seen across subjects. The extracted peaks included smaller peaks that other methods would generally ignore as noise. However, we chose to include them in the analysis.

### Statistical Model Fitting and Comparison

We fit four models to each subject's inter-pulse interval: the inverse Gaussian, lognormal, gamma, and exponential, each by computing maximum likelihood estimates of the parameters (34). We assessed goodness-of-fit by Akaike’s Information Criterion (AIC) and KS-plots. The AIC is defined as

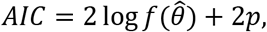

where 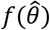 is the likelihood evaluated at the maximum likelihood estimate of the parameters and *p* is the number of parameters. A lower AIC indicates a better fit.

A KS-plot compares the model and empirical inter-pulse interval distributions using the time-rescaling theorem (35). The time-rescaling theorem states that any point process can be rescaled to a Poisson process with rate 1 using the conditional intensity function. The KS-distance computes the maximum distance between the quantiles of the rescaled data and the uniform distribution, which is a simple transform of a rate 1 Poisson process done for visual simplicity. A smaller KS distance indicates that the model is more similar to the empirical distribution. We computed a significance cut-off based on the number of extracted pulses and compared it to the KS-distances for each model (36). A KS-distance under the significance cutoff means that the model and empirical distributions are not significantly different from each other, whereas a KS-distance above the cutoff suggests that they are.

We also compared the models using a tail behavior analysis, in which the asymptotic settling rates of the models are compared to determine the heaviness of the tails of the distributions. We hypothesized that the models best fitting the EDA inter-pulse interval distribution were variations of the inverse Gaussian with slightly heavier or lighter tails (which can be captured by the lognormal and gamma distributions), rather than the exponential.

## Results

### Extraction of Galvanic Skin Response Events

Figure 1C shows an example of an excerpt of extracted pulses for one subject. This includes pulses large enough to be included in most analyses (traditional pulses) as well as those that are much smaller and usually either smoothed out or ignored as noise (smaller pulses). We included both types of pulses for all subjects and did not distinguish between them. The majority of subjects showed appreciable fluctuations in the tonic component, representing the background, across time (Figure S2). Across the 11 subjects, the total number of pulses in the one-hour time window ranged between 97 and 348, including the distantly spaced smaller pulses (Figure S3). The final prominence thresholds used ranged between 0.0025 and 0.023.

**Figure 1.**
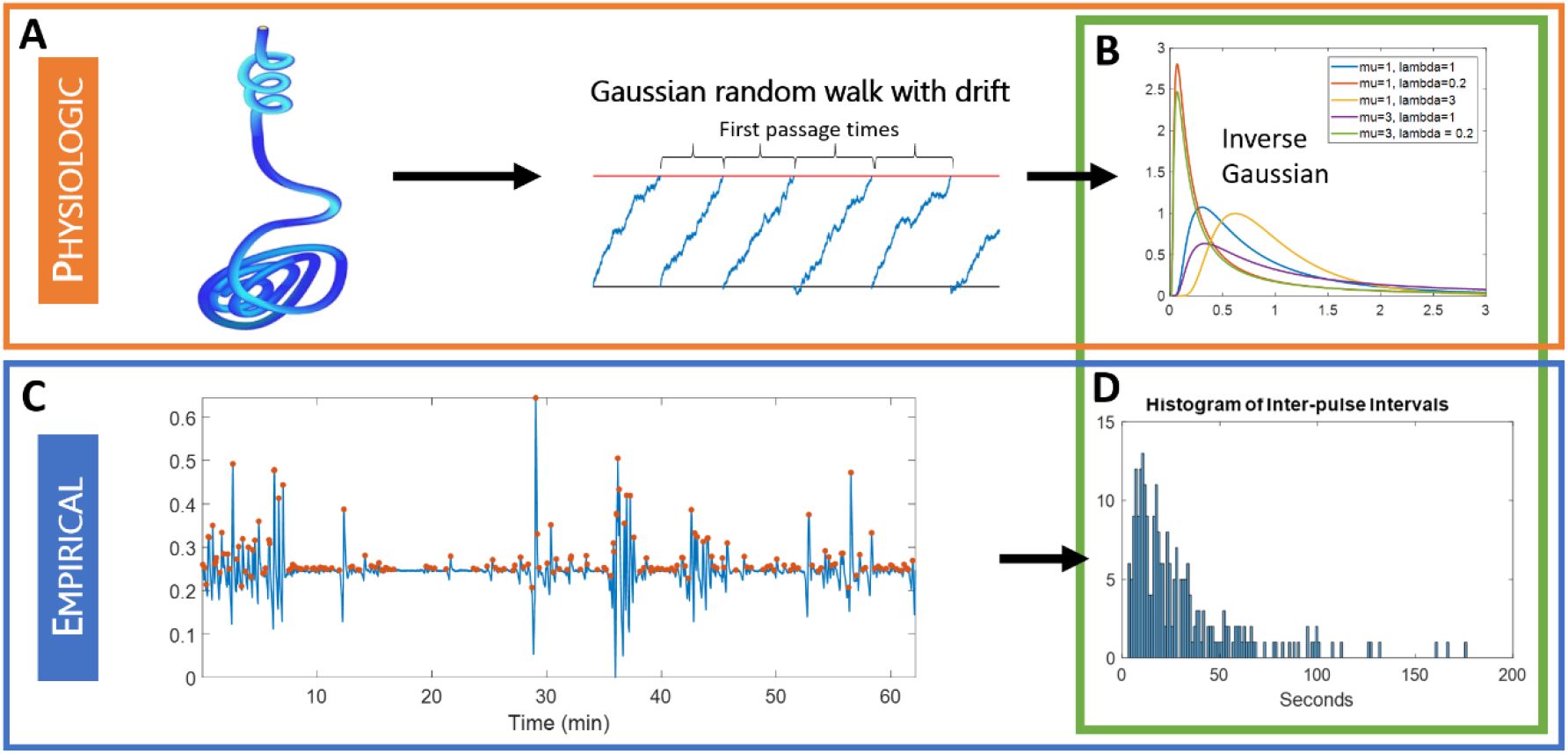
A summary of our model, including both physiologic and empirical components and how they align. (A) is an illustration of how sweat gland burst physiology can be modeled as a Gaussian random walk with drift, which suggests that the first passage times between events should follow (B) an inverse Gaussian distribution. (C) We extracted galvanic skin response events from EDA data, including both what are traditionally included as pulses in most analyses and small pulses, (D) and assessed the extent to which inter-pulse interval distributions are well described by inverse Gaussian, lognormal, gamma, and exponential probability models.

### Findings from Statistical Model Comparison

For all 11 subjects, one of the three models other than the exponential was the best fit model according to both AIC and KS-distance (Figures 2 and S4, Tables 1 and 2). Specifically, the exponential was the worst of the four models tested for 8 of the 11 subjects according to AIC and all 11 out of 11 subjects according to KS-distance. It is worthy of note that even if it was not the best fit model, the inverse Gaussian was a better fit than the exponential for 10 out of 11 subjects according to AIC and all 11 subjects according to KS-distance. Specifically, with respect to KS-distance, the inverse Gaussian and lognormal models were within the significance cutoff for all subjects, and the gamma for all but one, but the exponential was only within the cutoff for 4 out of 11 subjects. This suggests that for the majority of subjects, the exponential model was actually statistically significantly different from the data.

**Table 1.**
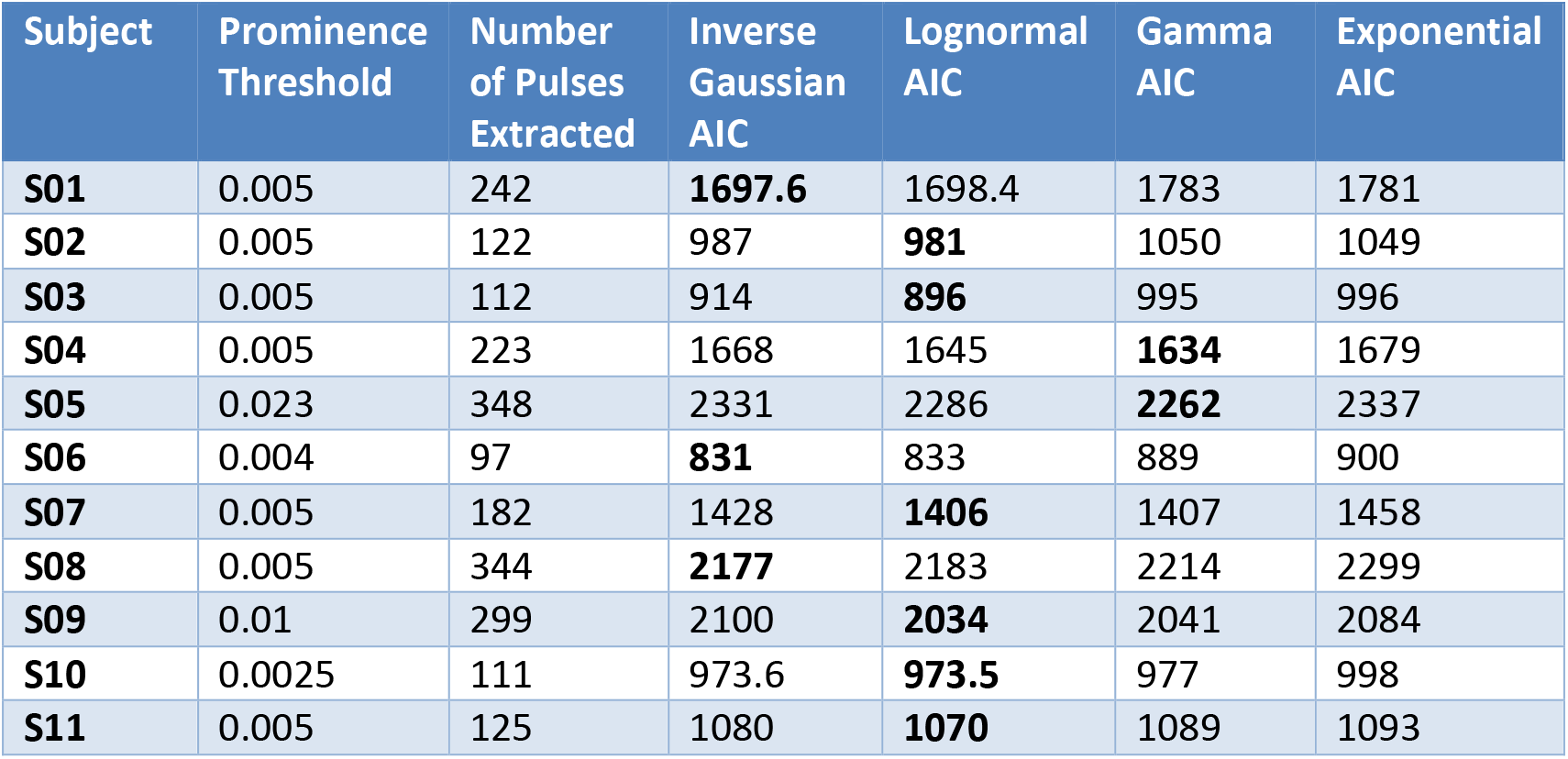
Results: AIC for all models (best model per subject in bold). The final prominence threshold used and number of pulses extracted is also indicated for each subject.

| Subject | Prominence Threshold | Number of Pulses Extracted | Inverse Gaussian AIC | Lognormal AIC | Gamma AIC | Exponential AIC |
| --- | --- | --- | --- | --- | --- | --- |
| **S01** | 0.005 | 242 | **1697.6** | 1698.4 | 1783 | 1781 |
| **S02** | 0.005 | 122 | 987 | **981** | 1050 | 1049 |
| **S03** | 0.005 | 112 | 914 | **896** | 995 | 996 |
| **S04** | 0.005 | 223 | 1668 | 1645 | **1634** | 1679 |
| **S05** | 0.023 | 348 | 2331 | 2286 | **2262** | 2337 |
| **S06** | 0.004 | 97 | **831** | 833 | 889 | 900 |
| **S07** | 0.005 | 182 | 1428 | **1406** | 1407 | 1458 |
| **S08** | 0.005 | 344 | **2177** | 2183 | 2214 | 2299 |
| **S09** | 0.01 | 299 | 2100 | **2034** | 2041 | 2084 |
| **S10** | 0.0025 | 111 | 973.6 | **973.5** | 977 | 998 |
| **S11** | 0.005 | 125 | 1080 | **1070** | 1089 | 1093 |

**Table 2.** Results: KS-distance for all models (best model per subject in bold, all models under significance cutoff marked with asterisk). The significance cut-off was computed based on the number of pulses extracted.

| Subject | Significance Cutoff | Inverse Gaussian KS-distance | Lognormal KS-distance | Gamma KS-distance | Exponential KS-distance |
| --- | --- | --- | --- | --- | --- |
| S01 | 0.087 | **0.057\*** | 0.065* | 0.079* | 0.085* |
| S02 | 0.123 | 0.073* | **0.040\*** | 0.119* | 0.120* |
| S03 | 0.128 | 0.121* | **0.054\*** | 0.159 | 0.169 |
| S04 | 0.091 | 0.077* | 0.043* | **0.023\*** | 0.102 |
| S05 | 0.073 | 0.068* | 0.031* | **0.030\*** | 0.122 |
| S06 | 0.138 | 0.071* | **0.050\*** | 0.134* | 0.177 |
| S07 | 0.101 | 0.088* | 0.057* | **0.048\*** | 0.146 |
| S08 | 0.073 | **0.025\*** | 0.030* | 0.047* | 0.120 |
| S09 | 0.079 | 0.078* | **0.028\*** | 0.051* | 0.100 |
| S10 | 0.129 | 0.050* | **0.042\*** | 0.055* | 0.109* |
| S11 | 0.122 | 0.052* | **0.028\*** | 0.056* | 0.077* |

**Figure 2.**
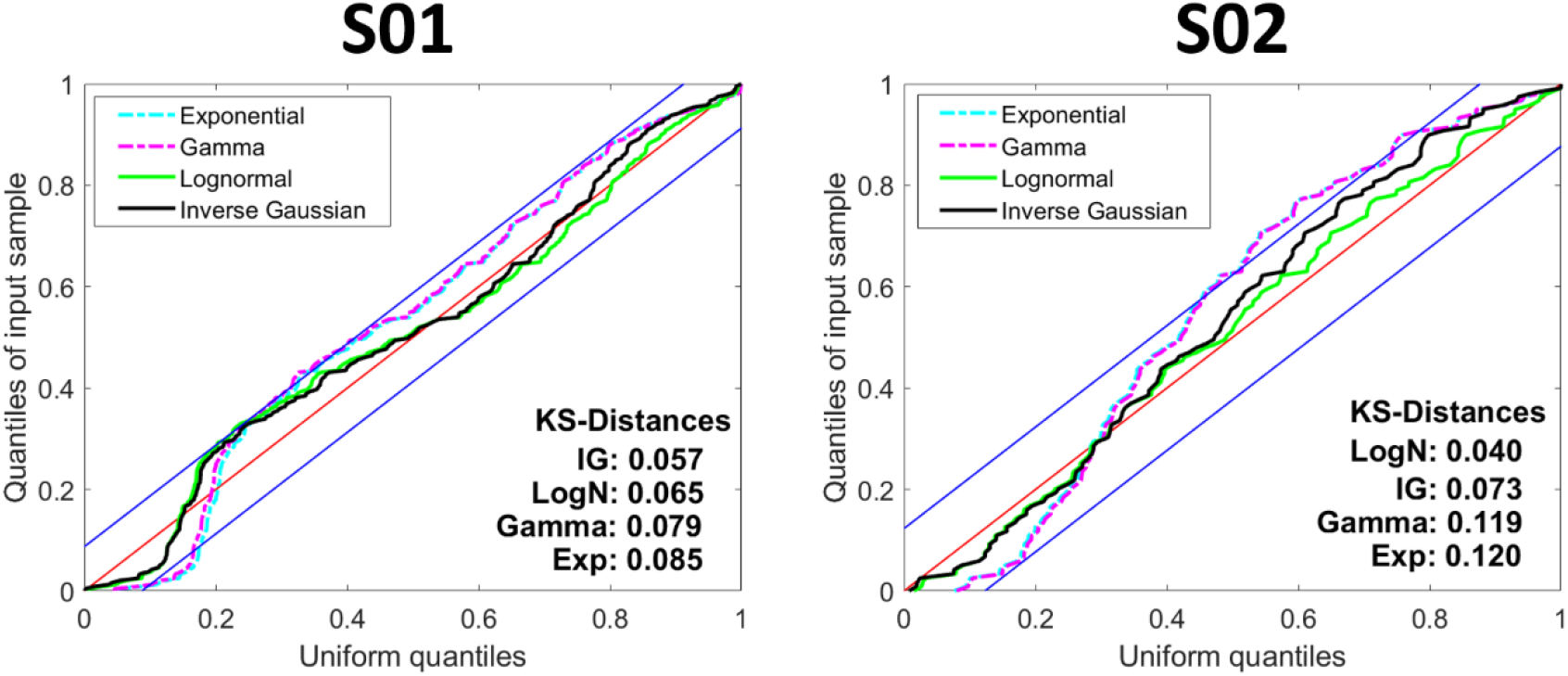
KS plots of inter-pulse interval data for Subjects S01 and S02 with 95% confidence bounds. All 4 models are shown against each other, with KS-distances for each. A smaller KS-distance, along with remaining fully within the 95% confidence bounds, indicates a better fit. The KS-distances are ordered in each case from best to worst fit model. ‘LogN’ refers to lognormal, ‘IG’ refers to inverse Gaussian, and ‘Exp’ refers to exponential. The inverse Gaussian is the best model for S01, whereas the lognormal for S02.

Both AIC and KS-distance were in agreement for the best fit model for 9 of the 11 subjects, with 2 subjects best fit by the inverse Gaussian (S01 and S08), 5 by the lognormal (S02, S03, S09, S10, and S11), and 2 by the gamma (S04 and S05). For S06 and S07, AIC and KS-distance returned different best fit distributions between inverse Gaussian and lognormal, and lognormal and gamma respectively. It is reasonable to expect that the results from AIC and from KS-distance will not match exactly since different metrics are intended to captured different aspects of model fits. However, the fact that they agree across the majority of subjects reinforces that there is in fact a clear difference between the models.

The second phase of comparing the models is using the tail behavior analysis (Table 3) which shows the settling rates predicted by all four models for all subjects. It is important to note that across all 11 subjects, the settling rate of the inverse Gaussian model is always less than that of the exponential, predicting a heavier tail than the exponential even though both are officially classified as medium-tailed distributions. Since the inverse Gaussian was also a better fit than the exponential for all 11 subjects (according to KS-distance), this supports our hypothesis that an inverse Gaussian model with a heavy tail not only accounts for the integrate-and-fire physiology, but also captures the smaller pulses occurring in regions of low background better than an exponential model. Both Subjects S01 and S08, for whom the inverse Gaussian was unanimously the best model, clearly demonstrate this result.

**Table 3.** Settling rates for all distributions; a lower settling rate indicates a heavier tail

|  | Inverse Gaussian | Lognormal | Gamma | Exponential |
| --- | --- | --- | --- | --- |
| Density | $(\frac{\lambda}{2\pi x^3})^{1/2} \exp(-\frac{\lambda(x-\mu)^2}{2\mu^2 x})$ | $(\frac{1}{2\pi\sigma^2})^{1/2} \exp\left(-\frac{(\ln x - \mu)^2}{2\sigma^2}\right) * \frac{1}{x}$ | $\frac{\beta^\alpha}{\Gamma(\alpha)} x^{\alpha-1} e^{-\beta x}$ | $\lambda e^{-\lambda x}$ |
| Settling Rate ($\tau_f$) | $\frac{\lambda}{2\mu^2}$ | 0 | $\beta$ | $\lambda$ |
| Tail Classification | Medium | Heavy | Medium | Medium |
| **S01** | $9.271/2(14.742)^2 = 0.0213$ | 0 | 0.071 | 0.068 |
| **S02** | $18.339/2(27.782)^2 = 0.0119$ | 0 | 0.035 | 0.036 |
| **S03** | $18.670/2(32.365)^2 = 0.0089$ | 0 | 0.026 | 0.031 |
| **S04** | $16.893/2(16.056)^2 = 0.0328$ | 0 | 0.120 | 0.062 |
| **S05** | $11.020/2(10.641)^2 = 0.0487$ | 0 | 0.184 | 0.094 |
| **S06** | $11.772/2(39.544)^2 = 0.0038$ | 0 | 0.017 | 0.025 |
| **S07** | $27.143/2(20.508)^2 = 0.0323$ | 0 | 0.107 | 0.049 |
| **S08** | $15.873/2(10.462)^2 = 0.0725$ | 0 | 0.198 | 0.096 |
| **S09** | $10.255/2(12.098)^2 = 0.0350$ | 0 | 0.142 | 0.083 |
| **S10** | $42.749/2(34.027)^2 = 0.0185$ | 0 | 0.056 | 0.029 |
| **S11** | $21.468/2(29.890)^2 = 0.0120$ | 0 | 0.044 | 0.033 |

The lognormal distribution by default has a heavier tail than the exponential and even than the inverse Gaussian, since it is officially classified as a heavy-tailed distribution (always has a settling rate of 0). It can also represent the combination of several inverse Gaussian models as the background or level of induced stimulation evolves over time. Therefore, Subjects S02, S03, S09, S10, and S11, for whom the lognormal was unanimously the best model, also align with our hypothesis about adequately capturing the sparse spontaneous pulses while accounting for integrate-and-fire physiology. Subject S06 was divided between the inverse Gaussian and lognormal as the best model, but the settling rate of the inverse Gaussian is very close to 0 (settling rate of the lognormal) for this subject compared to the settling rate of the exponential; therefore, this subject still fully aligns with our hypothesis.

Subjects S04 and S05 present an interesting case in which the gamma was the best model. Across all subjects, the gamma predicted a lighter tail than the inverse Gaussian model. However, for these two subjects, the asymptotic settling rate for the gamma model was much higher than even the exponential, which predicts a much lighter tail than the exponential. This can be explained by looking at the actual pulses extracted from the data (Figure S3). For both of these subjects, there are no long gaps between pulses; pulses are fairly constant throughout the dataset. Therefore, the inter-pulse interval distribution had almost no representation in the tail, which the gamma model is flexible enough to accommodate by making the tail very light. Subject S07 was divided between the lognormal and the gamma as the best model, each of which predicts a very different tail behavior. However, perhaps the tail was relatively underrepresented for this subject like for Subjects S04 and S05, so the fit was primarily based on the shape near the mode, which the lognormal model also fit well. With a data collection period longer than one hour, we could have captured more variation in dynamics.

## Discussion

In this study, we used EDA data collected from eleven healthy volunteers at rest to test our hypothesis that EDA contains highly regular statistical structure that is consistent with integrate-and-fire physiology that describes sweat gland function. To do that, we fit four different models to EDA, including the inverse Gaussian and exponential, and quantified the goodness-of-fit using two metrics, AIC and KS-distance. We also assessed tail behavior by settling rate. Together, we showed that the model fit and tail behavior are consistent with not only integrate-and-fire sweat gland physiology, but also the effect of varying EDA background on the dynamics of generated pulses.

The physiology of sweat gland activity predicts that the inter-pulse intervals for sweat gland pulses should be inverse Gaussian, which is an elementary model for processes where there is a gradual build-up of a continuous variable and a threshold has to be crossed to observe the event. Our results show that the data are more consistent with the inverse Gaussian model than the exponential, which represents Poisson activity for the pulses.

We further refined our hypothesis by taking into account that the background is gradually changing over time. This predicts that the inter-pulse intervals should follow a mixture of inverse Gaussian models, which can be modeled as gamma (lighter tail distributions) and lognormal (heavier tail distributions). More of the data are consistent with the lognormal heavier tail distribution, suggesting more frequent longer inter-pulse intervals than would be predicted by a homogeneous inverse Gaussian model. The fact that the inverse Gaussian still does very well reinforces the idea that the statistical structure in the data is fundamentally guided by the physiology of sweat gland activity which can be approximated well as an integrate-and-fire process (Gaussian random walk with drift).

This result creates a direct link between the physiology of sweat glands and the statistical structure of the data collected at the skin surface. The most detailed of existing models of EDA are founded in signal processing methods alone and require significant computational complexity. However, looking to the physiology provided a principled framework by which to drastically reduce dimensionality and increase the signal-to-noise ratio. This result has implications for understanding and tracking the sympathetic component of the autonomic nervous system in a more meaningful way.

In future work, we hope to use this result to inform an EDA analysis paradigm, which will allow us to ensure robust and accurate capture of the valuable statistical information contained in any EDA data set regardless of equipment used or condition studied. We will also study pulse amplitudes in more detail, taking into account that they depend both on stimulus amplitude and physiologic background state. Now that we have verified the presence of point process structure, we can study dynamics over time, applying history dependent inverse Gaussian models like those developed for heart rate variability (37–41). We will also study EDA in other contexts, such as during sleep, with pain, and under general anesthesia. Eventually, these methods have both clinical and non-clinical applications, such as in emotional state and stress detection. In all of these cases, because the underlying physiology is the same, the observed dynamics will likely provide distinct signatures for each of these important physiologic states.

## Acknowledgments

We would like to thank the MIT Clinical Research Center Staff. This work was partially funded by funds from the Picower Institute for Learning and Memory, the National Science Foundation Graduate Research Fellowship Program, and NIH Award P01-GM118629 (to E.N.B.).

