## Supplemental Figures for "Point Process Temporal Structure Characterizes Electrodermal Activity"

**This PDF file includes:**

Figures S1 to S4

**Additional Figures**

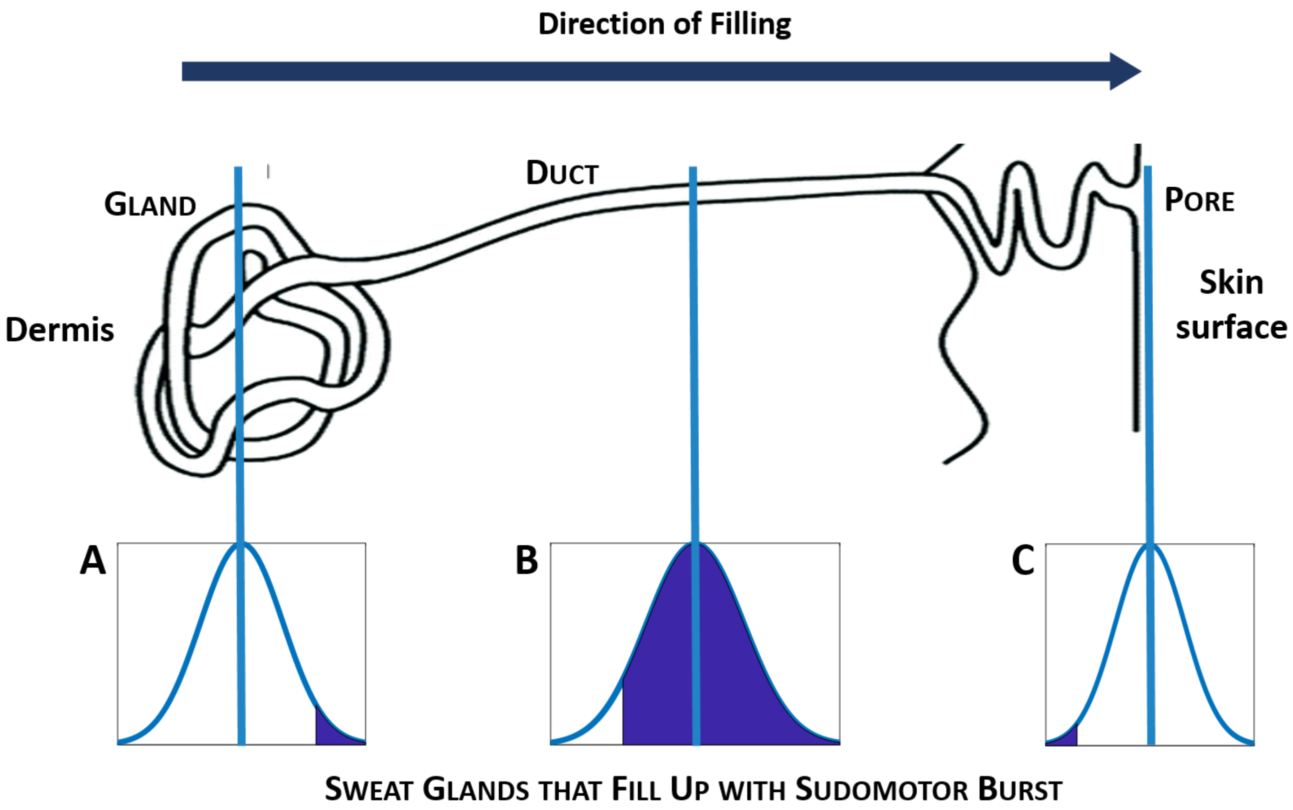


**Fig. S1.** A summary of our hypotheses around why the amplitudes of pulses are tied to underlying baseline physiological state. Different baseline or background levels represent different distributions of baseline filling of sweat glands. When a sudomotor burst results in sweat production, the sweat glands that contribute to the change in measured conductance are those that were not filled to the surface of the skin before but reach that point due to the sudomotor burst. The fraction of sweat glands for which this is likely to occur is determined by the baseline filling level. (A) When baseline filling is very low, only the most filled sweat glands will become fully filled with a sudomotor burst. This is a very small proportion, so pulse amplitude will be low. (B) When baseline filling is at some medium level, the large majority of glands will fill to the skin surface with a sudomotor burst, resulting in a large amplitude pulse. (C) Finally, when baseline filling is very high, only the lower tail of the distribution of glands will even be not already filled at baseline, which means that only these glands can contribute to a change in conductance with a sudomotor burst. This model helps explain the complex relationship between background and pulse amplitude.

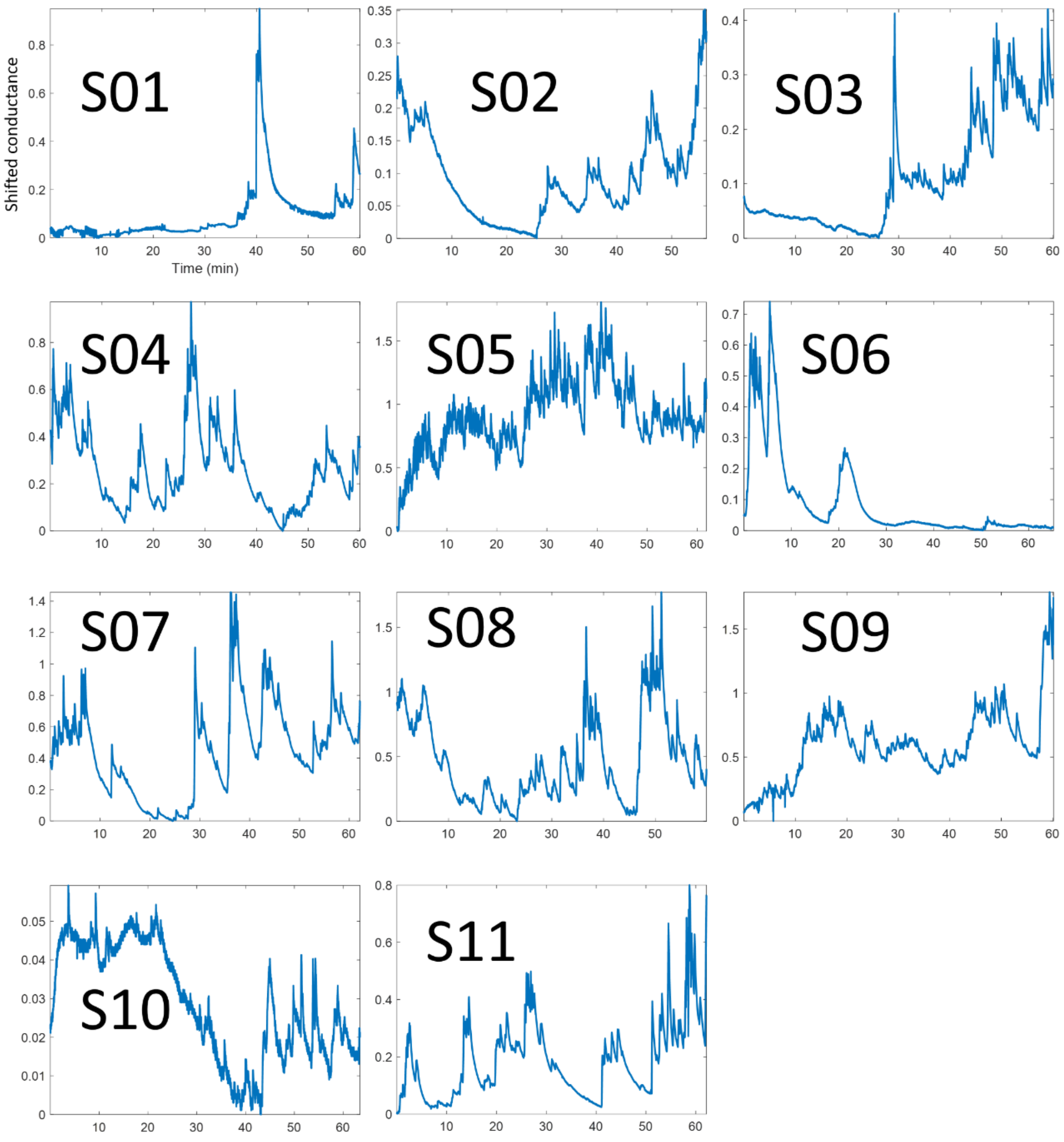


**Fig. S2.** The raw EDA data, with artifacts removed and minimum set to 0 for all subjects. This clearly shows the high degree of tonic (background) fluctuation for all subjects across time, as well as inter-subject heterogeneity in overall activity.

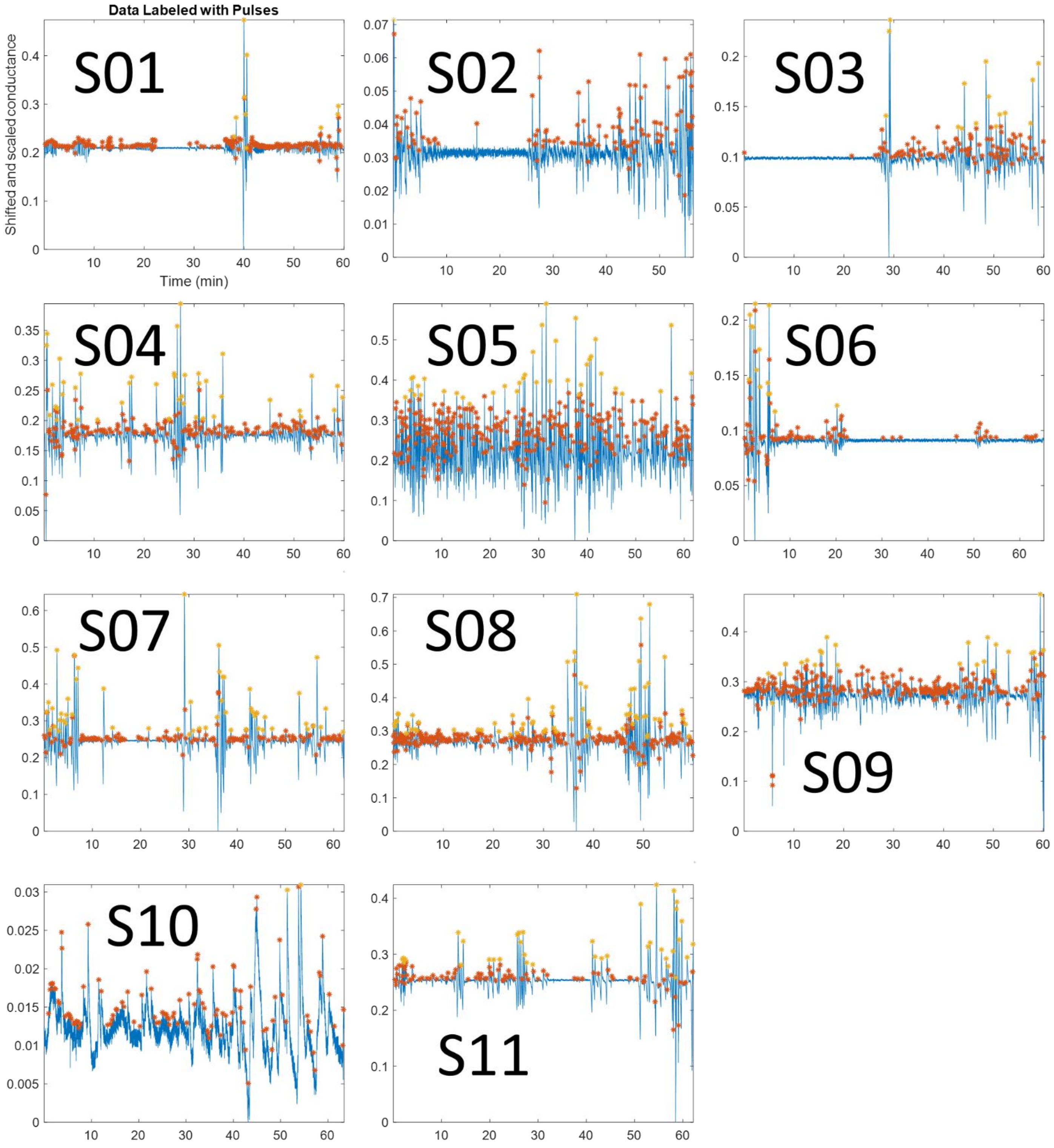


**Fig. S3.** The pulses extracted from the phasic component for all 11 subjects, with larger pulses labeled in yellow and smaller pulses in red. The data have already been preprocessed to remove artifacts and the tonic component.

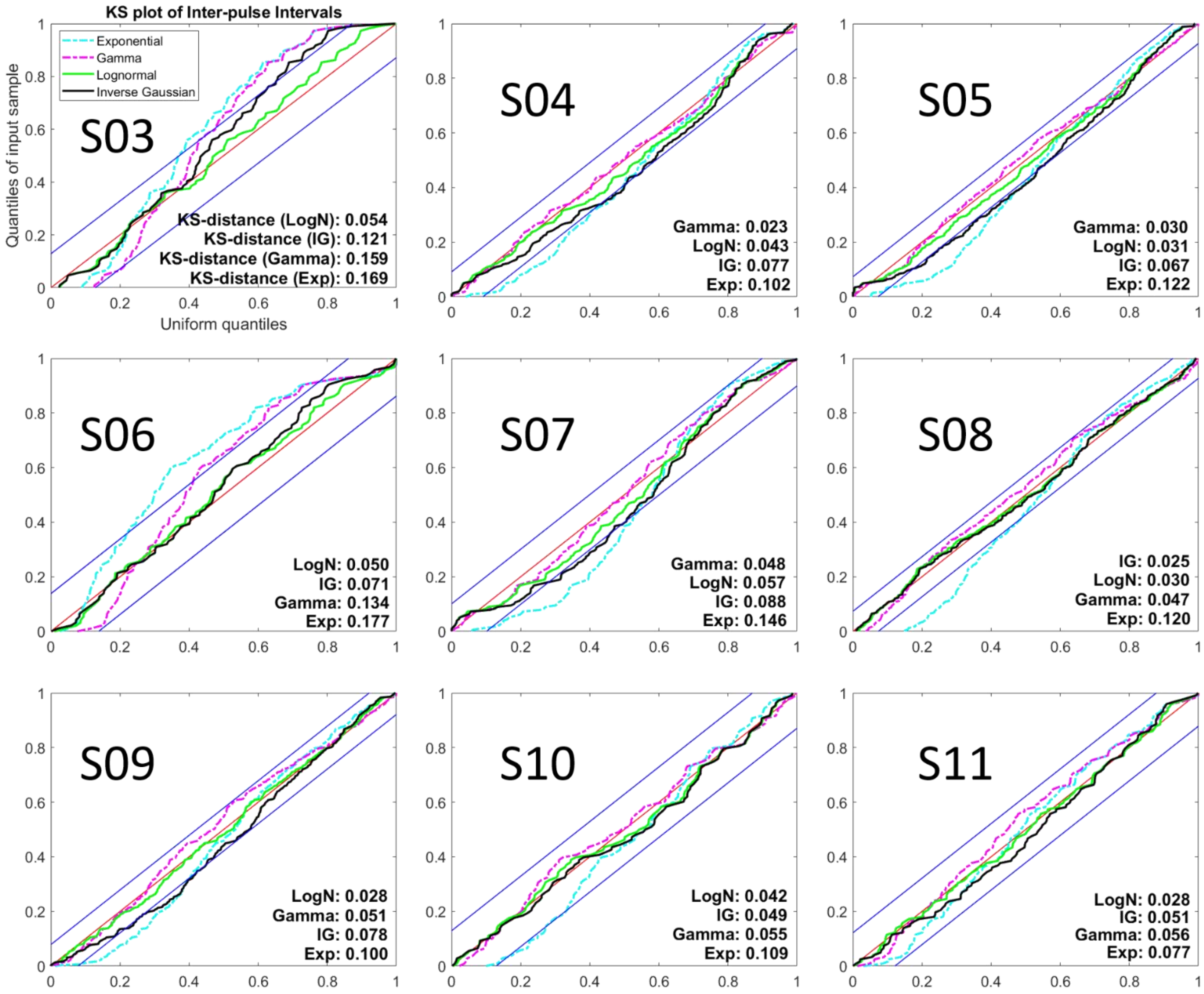


**Fig. S4.** KS plots with 95% confidence bounds for S03-S11. All 4 models are shown against each other, with KS-distances for each. A smaller KS-distance, along with remaining fully within the 95% confidence bounds, indicates a better fit. The KS-distances are ordered in each case from best to worst fit model.
